# Isthmin-1 is a Key Regulator of Induced Pluripotent Stem Cell–Derived Cardiomyocytes Maturation through Activation of p53 Signaling

**DOI:** 10.64898/2026.01.08.698535

**Authors:** Haowen Guo, Xin Zhou, Yang Shi, Bin Zhou, Jiaqi Tang, Faxiang Xu, Yanchen Guo, Fang Chen, Dongming Su, Qingguo Li

## Abstract

**Aims:** Human induced pluripotent stem cell–derived cardiomyocytes (iPSC-CMs) exhibit an immature structural and metabolic phenotype that limits their utility for cardiac disease modeling and regenerative therapy. Although multiple extrinsic strategies have been proposed to enhance iPSC-CM maturation, intrinsic molecular regulators governing this process remain incompletely defined. This study aimed to identify and characterize a novel molecular factor that promotes cardiomyocyte maturation.

**Methods and Results:** Cross-species transcriptomic analyses comparing adult versus fetal hearts and hypertrophic cardiomyopathy versus healthy myocardium identified Isthmin-1 (ISM1) as a conserved maturation-associated gene. In human iPSC-CMs, ISM1 overexpression enhanced sarcomere organization, mitochondrial oxidative phosphorylation, ATP production, and calcium-handling properties, whereas ISM1 knockdown impaired these maturation features. RNA sequencing revealed global transcriptional reprogramming toward an adult-like state, characterized by suppression of cell-cycle–related gene programs and activation of oxidative metabolic pathways. KEGG pathway enrichment, GSVA, and GSEA consistently identified p53 signaling as the most significantly activated pathway in ISM1-overexpressing iPSC-CMs. Mechanistically, ISM1 directly interacted with p53, enhanced its protein stability, promoted nuclear localization, and increased transcription of p53 downstream targets involved in metabolic remodeling and cell-cycle exit. Pharmacological inhibition of p53 abolished ISM1-induced structural and metabolic maturation, demonstrating that ISM1 promotes cardiomyocyte maturation in a p53-dependent manner.

**Conclusions:** ISM1 is a previously unrecognized molecular driver of cardiomyocyte maturation that promotes structural, metabolic, and functional maturation of iPSC-CMs through activation of p53-dependent transcriptional programs. These findings provide new mechanistic insight into cardiomyocyte maturation and identify ISM1 as a promising target for improving the maturity of stem cell–derived cardiomyocytes.

**Graphical Abstract:** 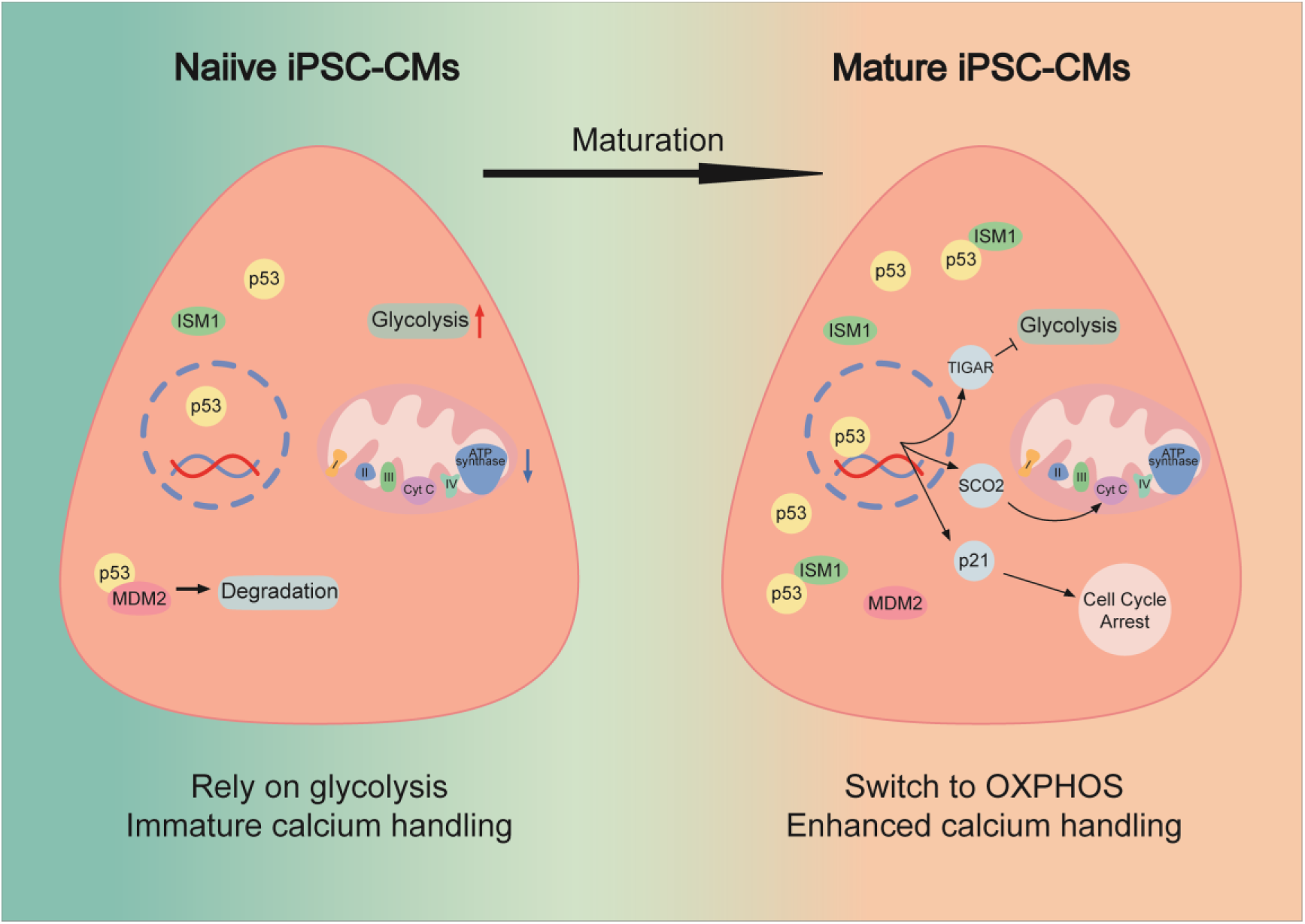

## Introduction

Human induced pluripotent stem cell–derived cardiomyocytes (iPSC-CMs) have emerged as a powerful platform for disease modeling, drug screening, and cardiac regenerative therapy. However, a major limitation of iPSC-CMs is their immature phenotype, which resembles fetal rather than adult cardiomyocytes. They exhibit disorganized sarcomeres, underdeveloped mitochondria, reliance on glycolytic metabolism, and immature calcium handling, all of which compromise contractile performance and electrical stability^1–3^. Enhancing the structural and metabolic maturation of iPSC-CMs is therefore essential to improve their physiological fidelity and clinical applicability. Despite extensive efforts using mechanical^4, 5^, biochemical^6–8^, and bioengineering strategies^9^, the molecular regulators that drive intrinsic cardiomyocyte maturation remain incompletely understood.

The need to promote iPSC-CM maturation parallels the necessity to maintain maturity in the adult heart, where pathological de-maturation has emerged as a key contributor to cardiac disease. In hypertrophic cardiomyopathy (HCM) and heart failure, mature cardiomyocytes revert to a fetal-like state, characterized by sarcomere disarray, metabolic reprogramming, and reactivation of fetal genes such as NPPA and NPPB^10–17^. This regression from oxidative to glycolytic metabolism, often termed a metabolic switch, undermines energy efficiency and contractile performance. Restoring or sustaining cardiomyocyte maturation—both in developing iPSC-CMs and in the diseased adult myocardium—therefore represents a convergent therapeutic strategy for heart failure prevention and cardiac regeneration.

In this study, we aimed to determine whether ISM1 functions as a “maturation factor” capable of promoting cardiomyocyte maturation. Using human iPSC-CMs, we demonstrate that ISM1 enhances sarcomere organization, mitochondrial oxidative phosphorylation, and calcium-handling function. Mechanistically, we identify that ISM1 exerts its effects through direct activation of p53 signaling, stabilizing and promoting the nuclear translocation of p53 to drive metabolic and transcriptional maturation.

Collectively, our findings define ISM1 as a dual-function regulator that promotes iPSC-CM maturation and potentially maintains adult cardiomyocyte maturity, bridging the developmental and pathological dimensions of cardiac remodeling. These insights provide a conceptual framework for targeting ISM1 to improve iPSC-CM–based transplantation therapies and prevent the progression of hypertrophic remodeling toward heart failure.

## Methods

### Cell culture and treatment

AC16 Cells were purchased from ZQXZbio and cultured in DMEM/F12 with 10% FBS. Human iPSC-derived cardiomyocytes (iPSC-CMs) were obtained from Nanjing HELP Therapeutics Co., Ltd and cultured in medium developed by Nanjing HELP Therapeutics Co., Ltd too. For ISM1 gain- and loss-of-function assays, iPSC-CMs were transduced with adenovirus expressing human ISM1 (Ad-ISM1) or transfected with siRNA targeting ISM1 (si-ISM1) using siRNA-Mate Plus (GenePharma). Neonatal rat cardiomyocytes (NRCMs) were isolated using the Pierce™ Primary Cardiomyocyte Isolation Kit and cultured in DMEM containing 10% FBS. To inhibit p53, cells were treated with Pifithrin-α (10 μM) for 48 h following ISM1 overexpression.

### Immunofluorescence

For immunofluorescence, cultured cardiomyocytes or heart sections were fixed, permeabilized with 0.1% Triton X-100, and incubated with primary antibodies against cTnT, p53, or ISM1 (Table S1), followed by fluorescence-conjugated secondary antibodies and DAPI nuclear staining. Images were captured using a Zeiss LSM 880 confocal microscope and analyzed with ImageJ software.

### Western blot analysis

Proteins were extracted in RIPA buffer containing protease and phosphatase inhibitors. Equal amounts of protein were resolved by SDS–PAGE, transferred to PVDF membranes, and incubated with specific primary and HRP-conjugated secondary antibodies. Signals were visualized using ECL detection (Tanon Imaging System) and quantified by ImageJ. β-actin or β-tubulin served as loading controls.

### Calcium handling assay

Intracellular calcium transients were recorded using Fluo-4 AM (5 μM) in spontaneously beating iPSC-CMs after 7 days of culture. Line-scan images were acquired with a Zeiss LSM 880 confocal microscope (488 nm excitation). Parameters including amplitude, upstroke and decay rates, and τ-decay were analyzed using GraphPad Prism 8.

### Seahorse extracellular flux assay

Mitochondrial respiration was measured using an Agilent Seahorse XFe96 Analyzer. iPSC-CMs were seeded on Matrigel-coated microplates and incubated in Seahorse XF Base Medium (10 mM glucose, 1 mM pyruvate, 2 mM glutamine, pH 7.4). Oxygen consumption rate (OCR) was recorded under basal conditions and after sequential injection of oligomycin (1.5 µM), FCCP (0.5–1.0 µM), and rotenone/antimycin A (0.5 µM each).

### Co-immunoprecipitation (Co-IP) and mass spectrometry

AC16 cells were lysed in IP buffer and incubated with anti-ISM1 or anti-p53 antibodies, followed by protein A/G magnetic beads. Immunoprecipitates were analyzed by Western blotting and LC–MS/MS to identify ISM1-binding partners. AlphaFold3 was used to simulate the ISM1–p53 complex interface.

### RNA-sequencing and bioinformatics analysis

RNA-seq was performed on control, ISM1-OE, and ISM1-KD iPSC-CMs (n = 3/group) using the Illumina NovaSeq 6000 platform. Reads were aligned to the human genome (GRCh38) using HISAT2, and DEGs were identified by DESeq2 (|log₂FC| > 0.5, FDR < 0.05). Functional enrichment was analyzed using ClusterProfiler for Gene Ontology (GO) and KEGG pathways.

### Statistical analysis

Data are presented as mean ± SEM. Statistical comparisons between two groups were made using unpaired two-tailed Student’s *t*-tests, and multiple groups were compared by one-way ANOVA with Tukey’s post hoc test. *P* < 0.05 was considered statistically significant.

## Results

### Identification and cross-species validation of maturation-associated genes in HCM

To explore the relationship between maturation and de-maturation processes during hypertrophic cardiomyopathy (HCM) pathogenesis, we analyzed differentially expressed genes (DEGs) from publicly available RNA-seq datasets comparing adult versus fetal hearts and HCM patients versus healthy controls (Figure 1A, B). The DEG profiles from these datasets were inversely correlated (Figure 1C), indicating that genes promoting maturation in the developing heart tend to be repressed during pathological de-maturation. Pathway enrichment analysis revealed that genes upregulated during maturation and downregulated in HCM were significantly enriched in fatty acid metabolism pathways (Figure S1A), consistent with the metabolic regression previously reported in HCM.

**Figure 1.**
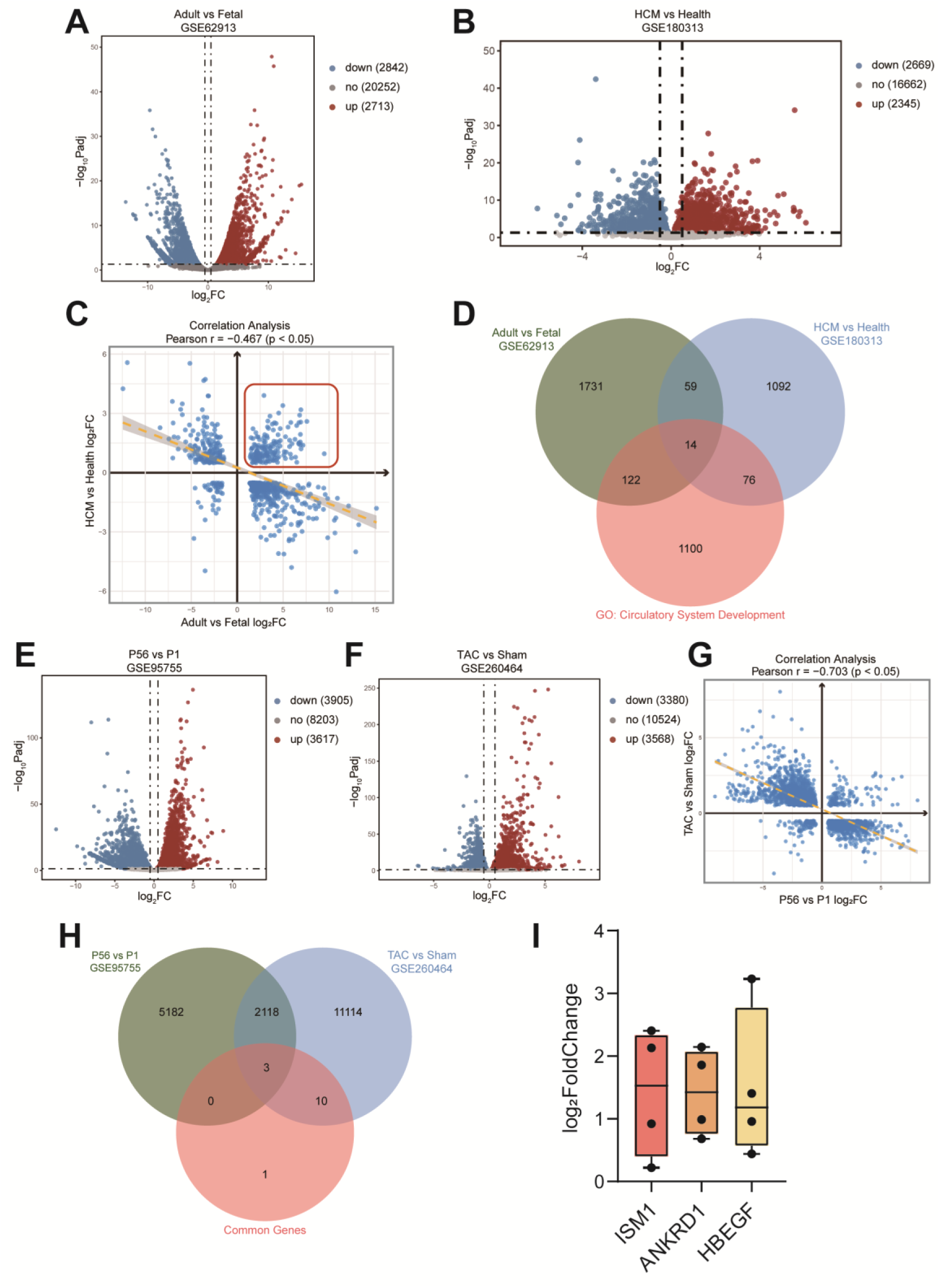
Identification and cross-species validation of maturation-associated genes in hypertrophic cardiomyopathy (HCM). (A, B) Volcano plots showing DEGs in adult versus fetal human hearts (GSE62913) and HCM versus healthy people (GSE180313). (C) Correlation analysis of DEGs between the two human datasets. (D) Fourteen shared upregulated genes identified in both human datasets under the GO term circulatory system development. (E, F) Volcano plots showing DEGs in mouse hearts comparing P56 versus P1 (GSE95755) and TAC versus sham conditions (GSE260464). (G) Correlation analysis of DEGs between the two mouse datasets. (H) Three shared upregulated genes in both mouse datasets compared with fourteen common genes identified in (D). (I) Log₂ fold-change comparison of the three conserved genes in four datasets.

Interestingly, a subset of genes was upregulated in both datasets, suggesting that these genes may represent compensatory or protective responses against pathological remodeling. To further refine these candidates, we intersected the commonly upregulated genes with those annotated under the Gene Ontology (GO) term circulatory system development, yielding 14 shared genes (Figure 1D).

To validate these findings across species, we next examined whether these genes exhibited conserved expression patterns in mice. Two RNA-seq datasets were analyzed: one comparing postnatal day 56 (P56) versus day 1 (P1) mouse hearts, representing developmental maturation, and another comparing transverse aortic constriction (TAC) versus sham hearts, representing stress-induced de-maturation (Figure 1E, F). Correlation analysis revealed consistent trends with the human datasets (Figure 1G). Genes upregulated during mouse heart maturation and downregulated after TAC were enriched in oxidative phosphorylation (OXPHOS) pathways (Figure S1B), further supporting the tight link between metabolic remodeling and cardiomyocyte maturity.

Among the 14 human candidate genes, three were commonly upregulated in both mouse datasets (Figure 1H). Comparative analysis of log₂ fold-change values identified Isthmin-1 (ISM1) as the most prominently upregulated gene (Figure 1I). Isthmin-1 (ISM1), originally identified as a secreted protein involved in embryonic development, angiogenesis, and metabolic regulation^18, 19^, has recently attracted attention for its potential role in tissue growth and stress adaptation. However, its function in the heart and cardiomyocyte maturation remains undefined. Through cross-species transcriptomic analyses, we identified ISM1 as one of the few genes upregulated during both developmental maturation and pathological hypertrophy, suggesting that it might act as a molecular link between maturation and de-maturation.

### ISM1 enhances structural, metabolic and functional maturation of induced pluripotent stem cell derived cardiomyocytes (iPSC-CMs)

To directly investigate the regulatory role of ISM1 during cardiomyocyte maturation, we utilized human induced pluripotent stem cell–derived cardiomyocytes (iPSC-CMs) as a well-established in vitro model that recapitulates many aspects of the immature cardiac phenotype. ISM1 was overexpressed in iPSC-CMs to evaluate its effects on structural organization, metabolic remodeling, and calcium-handling function, three key hallmarks of cardiomyocyte maturation.

Immunofluorescence staining of cTnT revealed well-organized, elongated sarcomeres in ISM1-overexpressing iPSC-CMs compared to control cells (Figure 2A, B). Immunoblot analysis confirmed a marked increase in cTnI protein abundance (Figure 2C, D), indicating enhanced assembly of contractile elements and structural maturation. ATP quantification showed significantly higher intracellular ATP levels in ISM1-overexpressing cells (Figure 2E), suggesting improved energy production capacity.

**Figure 2.**
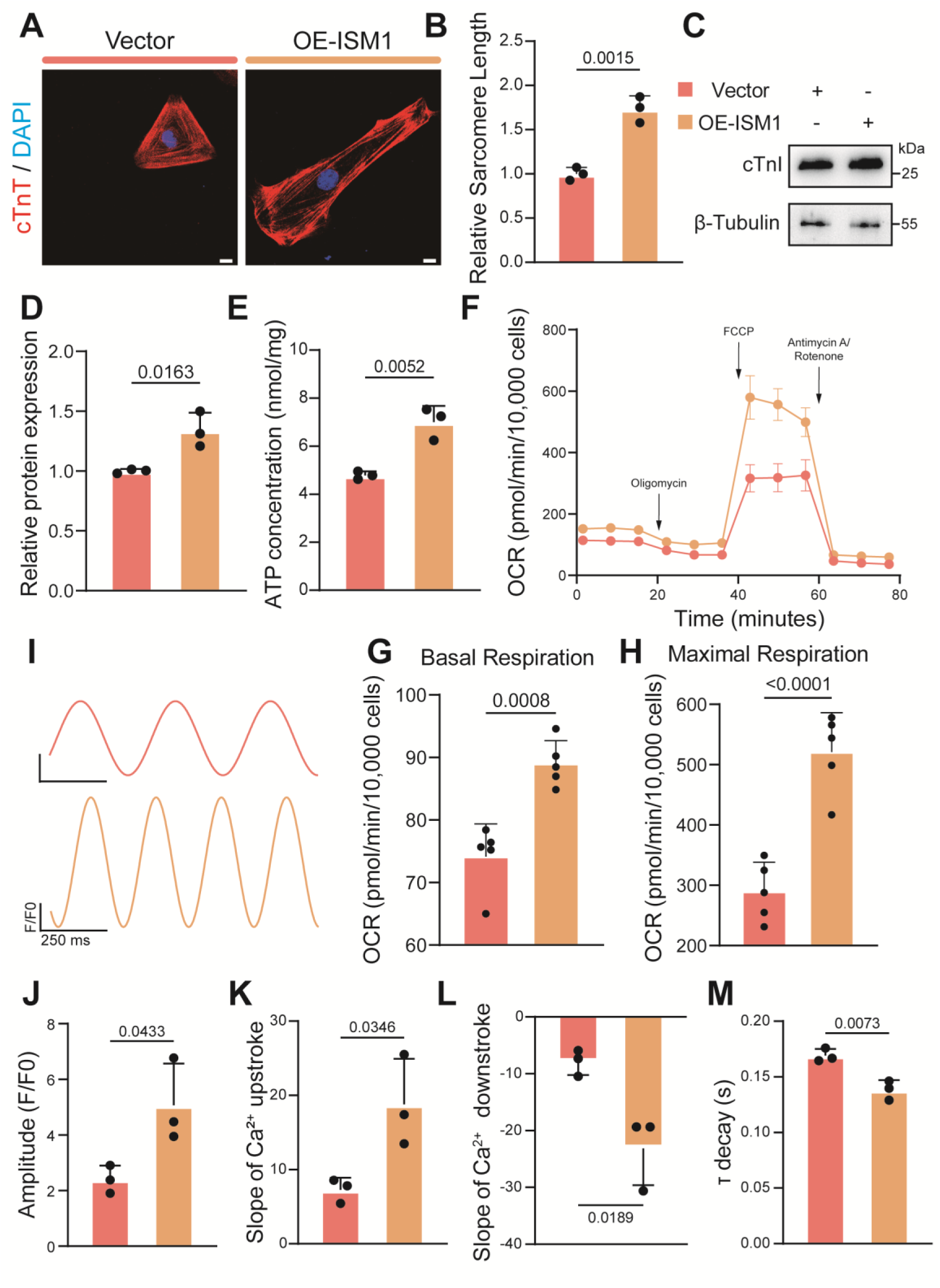
ISM1 promotes structural, metabolic, and functional maturation of induced pluripotent stem cell derived cardiomyocytes (iPSC-CMs). (A) Immunofluorescence staining of cTnT and DAPI in ISM1-overexpressing iPSC-CMs. Scale bar: 10 μm. (B) Quantification of sarcomere length (n=3). (C) Immunoblot analysis of cTnI expression in control and ISM1-overexpressing iPSC-CMs. (D) Quantification of cTnI protein levels normalized to β-tubulin (n=3). (E) Cellular ATP concentration in control and ISM1-overexpressing iPSC-CMs (n=3). (F–H) Seahorse analysis of oxygen consumption rate (OCR) and corresponding basal respiration and maximal respiration (n=5). (I) Representative Fluo-4 fluorescence recordings of calcium transients. (J–M) Quantification of calcium handling parameters, including amplitude, upstroke and downstroke slopes, and τ-decay time (n=3). Comparisons between two groups were performed using an unpaired two-tailed Student’s t-test, whereas one-way analysis of variance followed by Tukey post hoc test was conducted for comparisons among three or more groups. Values represent the mean ± SEM.

To further assess metabolic maturation, we performed Seahorse extracellular flux analysis to measure oxidative metabolism. ISM1 overexpression markedly increased basal respiration, maximal respiration, and ATP-linked respiration (Figure 2F–H), demonstrating a metabolic shift from glycolysis toward oxidative phosphorylation (OXPHOS)—a defining feature of mature cardiomyocytes. These results indicate that ISM1 not only enhances mitochondrial function but also promotes energy efficiency and metabolic specialization during maturation. In terms of functional maturation, Fluo-4 calcium imaging revealed significantly improved calcium handling (Figure 2I). ISM1-overexpressing iPSC-CMs exhibited higher calcium transient amplitude, faster upstroke and decay kinetics, and prolonged τ-decay time (Figure 2J–M), all reflecting a more synchronized excitation–contraction coupling typical of adult cardiomyocytes.

Together, these findings demonstrate that ISM1 facilitates structural, metabolic, and functional maturation of iPSC-CMs. By promoting sarcomere organization, enhancing mitochondrial oxidative metabolism, and refining calcium-handling properties, ISM1 emerges as a key regulator capable of advancing iPSC-CMs toward a mature, energetically competent, and physiologically functional state—paralleling its role in maintaining cardiac maturity in vivo.

### ISM1 is required for structural, metabolic, and functional maturation of induced pluripotent stem cell derived cardiomyocytes (iPSC-CMs)

To determine whether ISM1 is indispensable for cardiomyocyte maturation, we silenced ISM1 expression in iPSC-CMs using siRNA-mediated knockdown. Loss of ISM1 led to pronounced immaturity in both cellular morphology and metabolic function. Immunofluorescence staining of cTnT revealed shortened and disorganized sarcomeres in ISM1-deficient cells compared with controls (Figure 3A, B), and immunoblotting of cTnI confirmed a significant reduction in contractile protein expression (Figure 3C, D), indicating impaired structural maturation.

**Figure 3.**
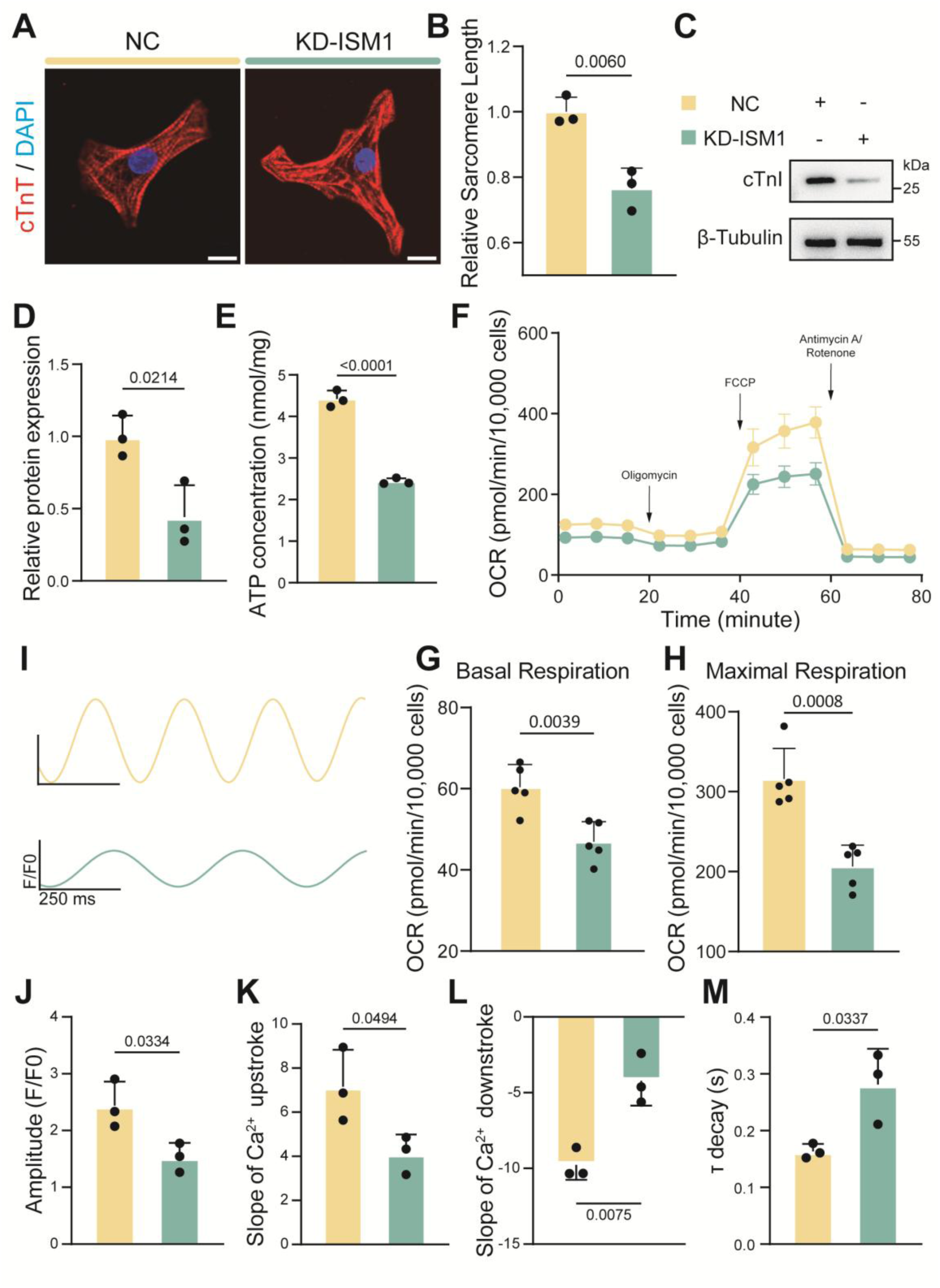
Silencing ISM1 impairs structural, metabolic, and functional maturation of induced pluripotent stem cell derived cardiomyocytes (iPSC-CMs). (A) Immunofluorescence staining of cTnT in ISM1-knockdown iPSC-CMs. Scale bar: 10 μm. (B) Quantification of sarcomere length (n=3). (C) Western blot analysis of cTnI expression in control (NC) and ISM1-knockdown (KD) iPSC-CMs. (D) Quantification of cTnI protein levels normalized to β-tubulin (n=3). (E) Cellular ATP concentration in control (NC) and ISM1-knockdown (KD) iPSC-CMs (n=3). (F–H) Seahorse analysis of oxygen consumption rate (OCR) and corresponding basal respiration and maximal respiration (n=5). (I) Representative Fluo-4 recordings of calcium transients. (J–M) Quantification of calcium handling parameters, including amplitude, upstroke and downstroke slopes, and τ-decay time (n=3). Comparisons between two groups were performed using an unpaired two-tailed Student’s t-test, whereas one-way analysis of variance followed by Tukey post hoc test was conducted for comparisons among three or more groups. Values represent the mean ± SEM.

At the metabolic level, ATP quantification showed a notable decline in intracellular ATP content in ISM1-deficient iPSC-CMs (Figure 3E), reflecting reduced energy production capacity. Seahorse extracellular flux analysis further demonstrated diminished oxidative metabolism, characterized by lower basal and maximal oxygen consumption rates and reduced ATP-linked respiration (Figure 3F–H). These results suggest that ISM1 knockdown hinders the metabolic transition from glycolysis to oxidative phosphorylation (OXPHOS) — a defining hallmark of cardiomyocyte maturation.

Functional assessments using Fluo-4 calcium imaging revealed clear defects in excitation–contraction coupling. ISM1-deficient iPSC-CMs exhibited reduced calcium transient amplitude, slower upstroke and downstroke kinetics, and an accelerated τ-decay (Figure 3I–M), indicating impaired calcium cycling and delayed electrophysiological maturation.

Together, these findings establish that ISM1 is essential for cardiomyocyte maturation. Its absence disrupts sarcomere organization, mitochondrial metabolic activation, and calcium-handling function, leading to a globally immature phenotype. Consistent with our overexpression results, these data highlight ISM1 as a core regulator of cardiomyocyte structural and metabolic development, necessary for achieving and sustaining the mature, energetically efficient state characteristic of adult cardiac cells.

### ISM1 Reprograms the Transcriptional Landscape of iPSC-CMs to Drive Structural, Metabolic, and Cell-Cycle Maturation

To elucidate the molecular mechanisms through which ISM1 promotes cardiomyocyte maturation, we performed RNA-sequencing (RNA-seq) on iPSC-CMs with ISM1 overexpression (OE), ISM1 knockdown (KD), and corresponding controls (n = 3 per group). Principal component analysis (PCA) demonstrated clear separation among the three conditions (Figure S2A), indicating robust ISM1-dependent transcriptional reprogramming.

Differential expression analysis identified 5,773 upregulated and 5,849 downregulated genes in ISM1-overexpressing iPSC-CMs relative to controls (FDR < 0.05) (Figure 4A). Key maturation-associated genes, including *MYH6* and *KCNJ2*, were significantly upregulated, whereas immature or fetal genes (*CDK1, NPPA, NPPB*) were downregulated, consistent with the enhanced structural and metabolic maturation observed experimentally. Conversely, ISM1 knockdown resulted in 1,840 upregulated and 2,053 downregulated genes (Figure S2B), with canonical maturation markers such as *ATP2A2, TNNT2, CPT1B*, and *HCN4* significantly decreased—reflecting impaired maturation phenotypes.

**Figure 4.**
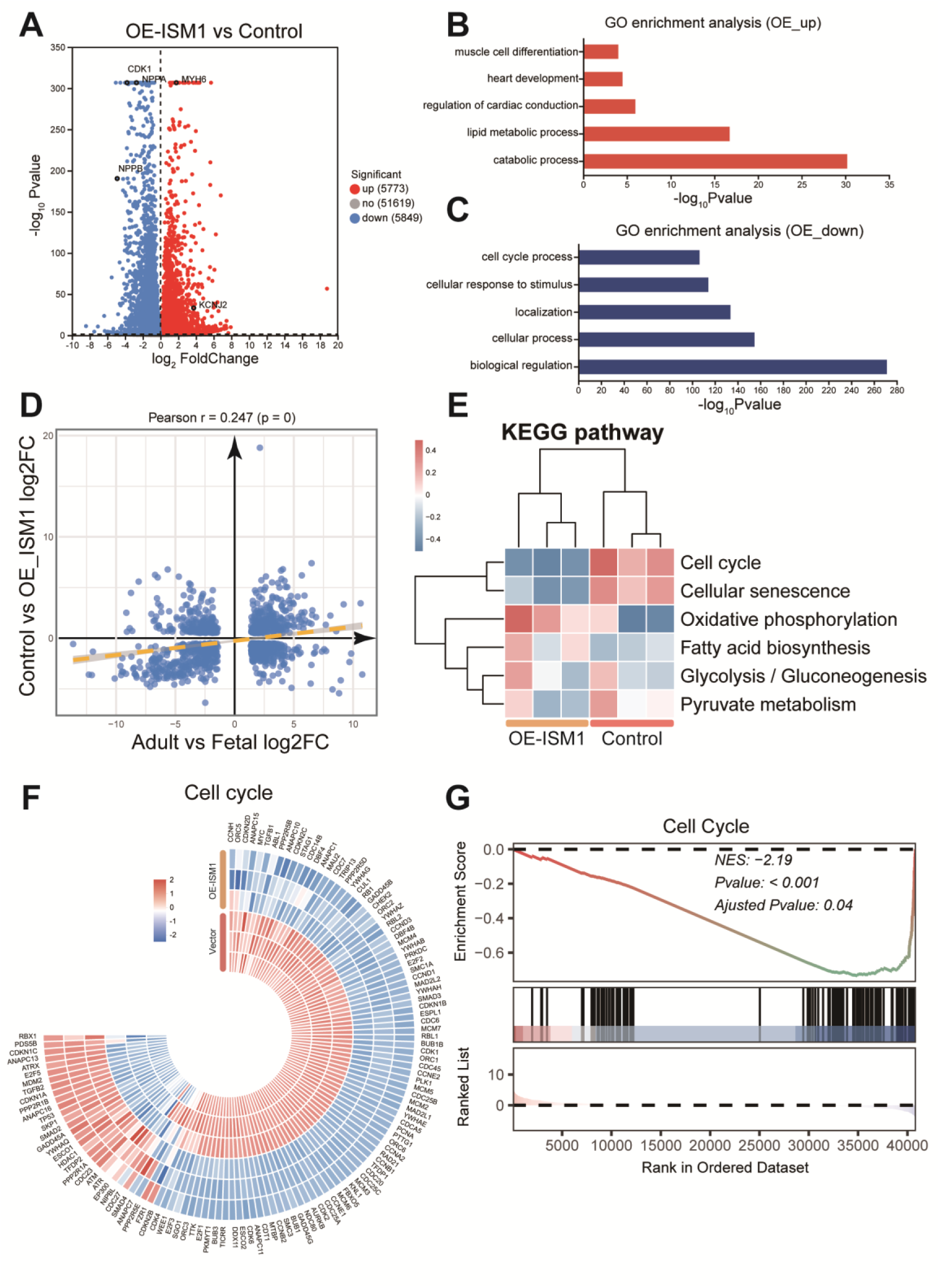
ISM1 Reprograms the Transcriptional Landscape of iPSC-CMs to Drive Structural, Metabolic, and Cell-Cycle Maturation. (A) Volcano plot showing differentially expressed genes (DEGs) in OE versus control. (B, C) Gene Ontology (GO) enrichment of upregulated and downregulated DEGs in OE versus control. (D) Correlation analysis of DEGs between ISM1-OE iPSC-CMs and adult-versus-fetal human heart datasets. (E) Gene set variation analysis (GSVA) of targeted KEGG pathways in ISM1-overexpressing iPSC-CMs. (F) Heatmap of DEGs in the cell cycle pathway in ISM1-overexpressing iPSC-CMs. (G) Gene Set Enrichment Analysis (GSEA) reveals suppression of the cell cycle pathway in ISM1-overexpressing iPSC-CMs. Comparisons between two groups were performed using an unpaired two-tailed Student’s t-test, whereas one-way analysis of variance followed by Tukey post hoc test was conducted for comparisons among three or more groups. Values represent the mean ± SEM.

Gene Ontology (GO) enrichment analysis revealed that OE-upregulated genes were enriched in catabolic and lipid metabolic processes, while downregulated genes were involved in cell-cycle regulation and biological control (Figure 4B, C), indicating a transcriptional shift from proliferation toward maturation. In contrast, KD-upregulated genes were enriched in negative regulation of cellular and metabolic processes, and KD-downregulated genes clustered in developmental and metabolic pathways (Figure S2C, S2D), demonstrating that ISM1 loss disrupts mature gene expression programs and promotes transcriptional regression. Notably, DEG profiles from ISM1-OE iPSC-CMs positively correlated with adult-versus-fetal human heart datasets (Figure 4D), further supporting that ISM1 shifts iPSC-CMs toward an adult-like molecular phenotype.

To further characterize pathway-level changes, we performed GSVA on KEGG maturation-related pathways (Figure 4E). ISM1 overexpression led to a marked downregulation of cell-cycle activity and upregulation of oxidative phosphorylation (OXPHOS). Most cell-cycle–related genes were significantly suppressed (Figure 4F), and GSEA confirmed strong enrichment of cell-cycle inhibition (Figure 4G). In parallel, a predominant upregulation of OXPHOS genes (Figure S2E) supports the conclusion that ISM1 facilitates cell-cycle exit and metabolic transition from glycolysis to oxidative metabolism, thereby promoting iPSC-CM maturation.

### Direct ISM1–p53 Interaction Underlies Activation of Maturation-Associated Transcriptional Pathways

To investigate how ISM1 reprograms the transcriptional landscape of iPSC-CMs, we first performed KEGG pathway enrichment analysis, which identified the p53 signaling pathway as the most significantly enriched pathway in ISM1-overexpressing cells (Figure 5A). The p53 pathway is a central regulator of mitochondrial biogenesis, oxidative phosphorylation, glycolytic suppression, and cell-cycle exit—all fundamental components of cardiomyocyte maturation. Consistent with this, key p53-associated genes, including *TP53, CDKN1A, PPARGC1A, TIGAR,* and *SCO2*, were markedly upregulated in ISM1-overexpressing iPSC-CMs (Figure S3A), indicating activation of p53-mediated metabolic remodeling and cell-cycle silencing.

**Figure 5.**
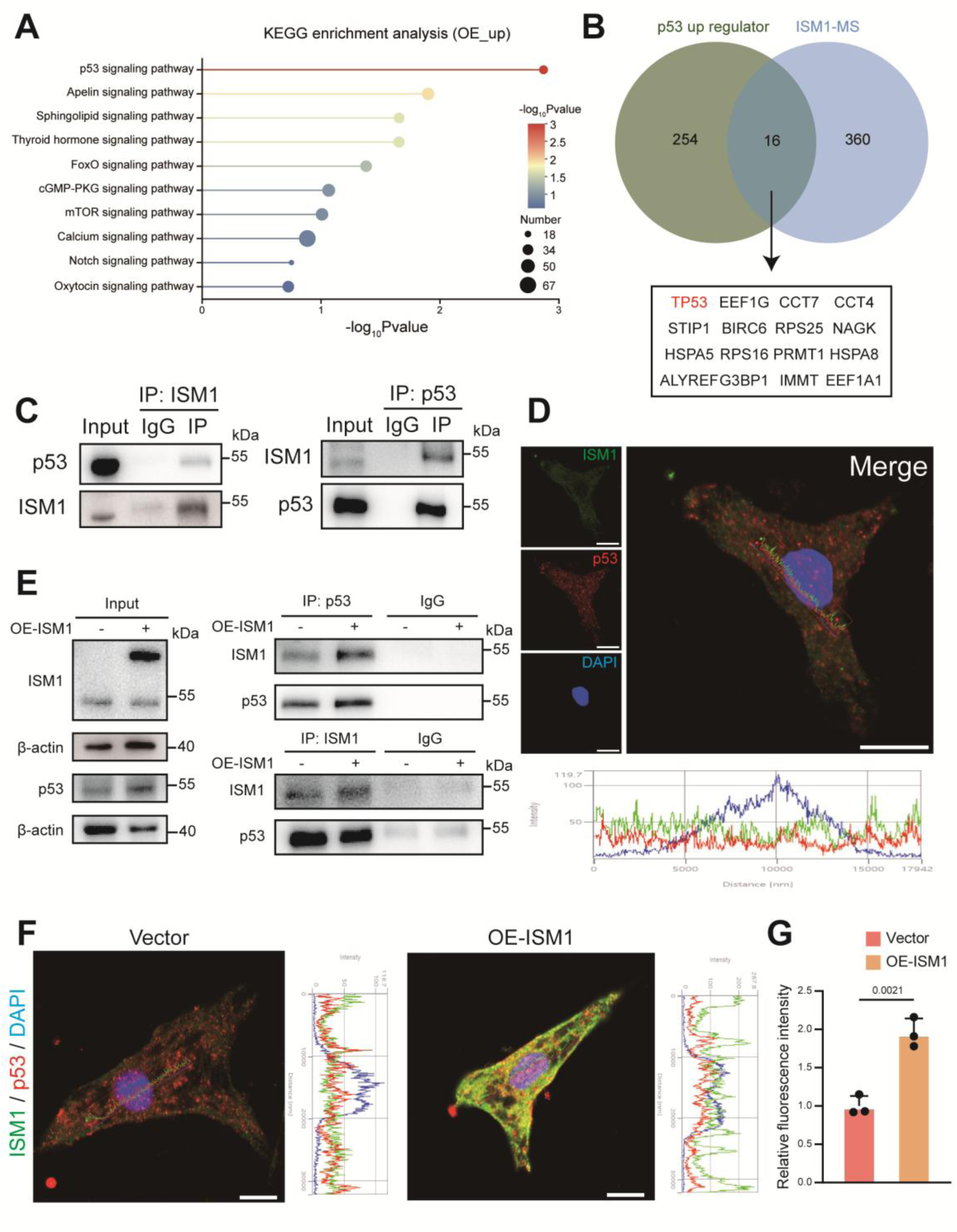
Direct ISM1–p53 Interaction Underlies Activation of Maturation-Associated Transcriptional Pathways. (A) KEGG pathway enrichment analysis in OE iPSC-CMs. (B) Co-immunoprecipitation and mass spectrometry (Co-IP/MS) analysis showing ISM1-binding proteins, including p53 and 15 other p53-upregulator. (C) Co-immunoprecipitation (Co-IP) and Western blot (WB) confirming reciprocal binding between ISM1 and p53 in AC16 cells. (D) Immunofluorescence (IF) of ISM1 and p53 in NRCMs. Scale bar: 10 μm. (E) Co-immunoprecipitation (Co-IP) and Western blot (WB) of ISM1 and p53 after overexpressing ISM1 in AC16 cells. (F) Immunofluorescence (IF) of ISM1 and p53 in ISM1 overexpressing NRCMs. Scale bar: 10 μm. (G) Quantification of p53 in ISM1 overexpressing NRCMs (n=3). Comparisons between two groups were performed using an unpaired two-tailed Student’s t-test, whereas one-way analysis of variance followed by Tukey post hoc test was conducted for comparisons among three or more groups. Values represent the mean ± SEM.

To further elucidate how ISM1 influences p53 signaling, we performed co-immunoprecipitation coupled with mass spectrometry (Co-IP/MS). This analysis identified 376 ISM1-interacting proteins, including 16 known upstream activators of the p53 pathway (Figure 5B). Notably, p53 itself was detected among ISM1-binding partners, suggesting that ISM1 may directly modulate p53 transcriptional activity and protein stability, thereby promoting maturation.

We next validated this interaction using Co-IP assays in AC16 cells. ISM1 immunoprecipitates contained a distinct p53 band, and reciprocal pulldown with anti-p53 antibody recovered the ISM1 protein, with no detectable signal in IgG controls (Figure 5C). Immunofluorescence staining further revealed co-localization of ISM1 and p53 (Figure 5D), supporting a direct association in situ.

Upon ISM1 overexpression, the amount of ISM1 bound to p53 significantly increased (Figure 5E), although the total ISM1 detected in p53 pulldowns remained similar, likely due to the higher endogenous abundance of p53. IF imaging also showed enhanced co-localization of ISM1 and p53 following ISM1 overexpression (Figure 5F). In addition, total p53 protein levels were elevated in ISM1-overexpressing cells (Figure 5G, S3B), suggesting that ISM1 may stabilize p53 or reduce its degradation.

### ISM1 Enhances p53 Stability, Nuclear Activity, and Downstream Transcription to Drive Maturation

To determine whether ISM1 regulates p53 protein stability, we performed cycloheximide (CHX) chase assays, which revealed that ISM1 significantly delayed p53 degradation and prolonged its half-life (Figure 6A, B). To assess whether stabilized p53 retains its functional capacity, we conducted nuclear–cytoplasmic fractionation, which showed a marked increase in nuclear p53 levels following ISM1 overexpression (Figure 6C). Consistently, luciferase reporter assays using p53-responsive promoters demonstrated enhanced p53 transcriptional activity and elevated expression of downstream targets (*CDKN1A, TIGAR, SCO2*) in ISM1-overexpressing cells (Figure 6D–F). These findings collectively demonstrate that ISM1 directly binds to p53, enhances its stability, promotes nuclear translocation, and activates p53 signaling, thereby driving transcriptional and metabolic programs essential for iPSC-CM maturation.

**Figure 6.**
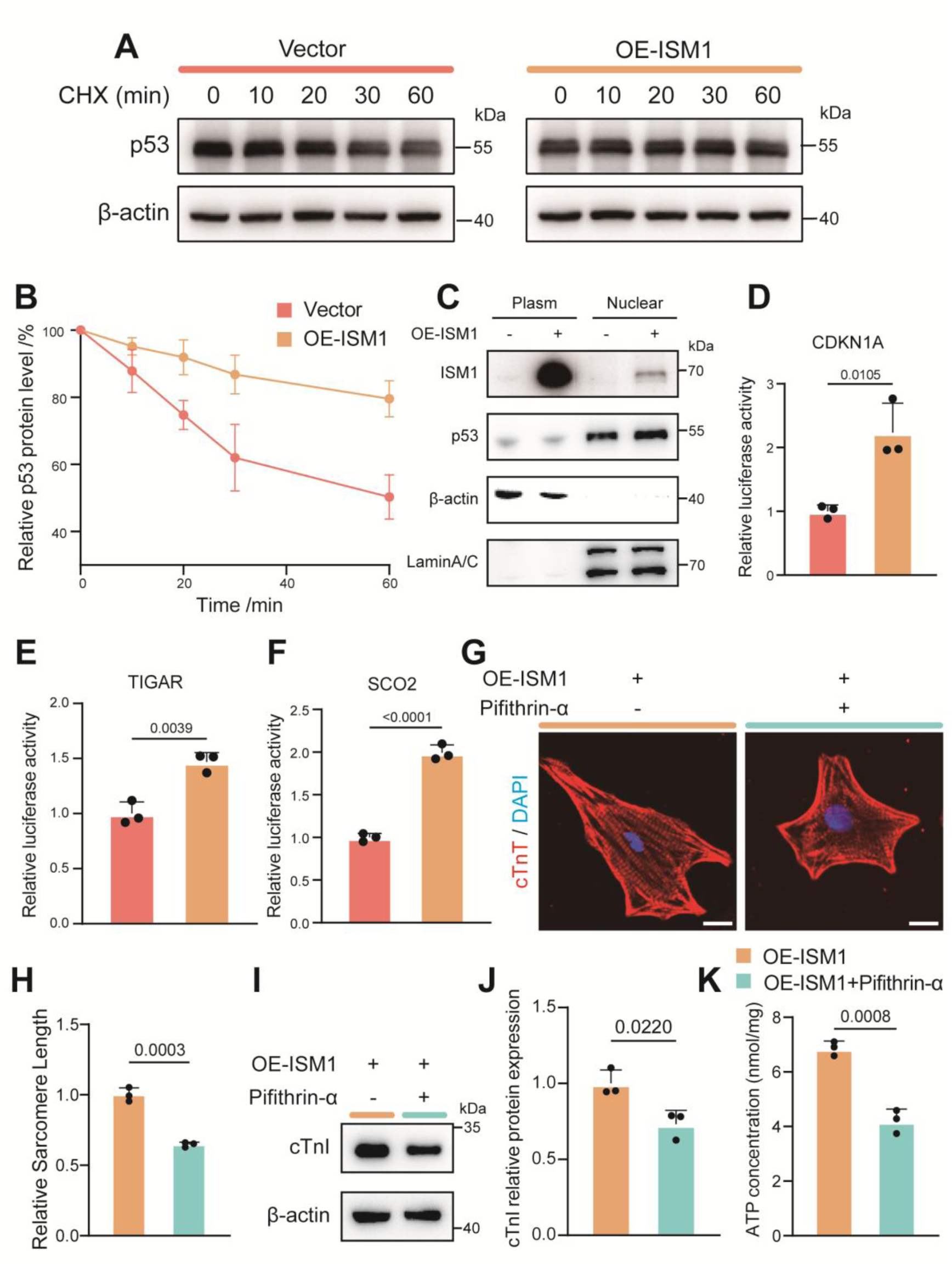
ISM1 Enhances p53 Stability, Nuclear Activity, and Downstream Transcription to Drive Maturation. (A, B) Cycloheximide (CHX) chase assay of p53 in ISM1-overexpressing AC16 cells (n=3). (C) Co-immunoprecipitation and mass spectrometry (Co-IP/MS) analysis showing ISM1-binding proteins, including p53 and 15 other p53-upregulator. (C) Western blot (WB) of nuclear–cytoplasmic fractionation. (D-F) Luciferase reporter assays for p53 target genes (*CDKN1A*, *TIGAR*, *SCO2*) in AC16 cells (n=3). (G) Immunofluorescence staining of cTnT in ISM1-overexpression and p53-inhibition iPSC-CMs. Scale bar: 10 μm. (H) Quantification of sarcomere length (n=3). (I) Western blot analysis of cTnI expression in ISM1-overexpression and p53-inhibition iPSC-CMs. (J) Quantification of cTnI protein levels normalized to β-actin (n=3). (K) Cellular ATP concentration in ISM1-overexpression and p53-inhibition iPSC-CMs (n=3). Comparisons between two groups were performed using an unpaired two-tailed Student’s t-test, whereas one-way analysis of variance followed by Tukey post hoc test was conducted for comparisons among three or more groups. Values represent the mean ± SEM.

To confirm that ISM1’s function depends on p53 activity, we treated ISM1-overexpressing iPSC-CMs with Pifithrin-α, a pharmacological inhibitor of p53. p53 inhibition abrogated the maturation-promoting effects of ISM1: immunofluorescence staining of cTnT revealed shorter and disorganized sarcomeres (Figure 6G, H), and immunoblotting of cTnI showed a pronounced reduction in contractile protein expression (Figure 6I, J), indicating impaired structural maturation. Metabolically, ATP quantification demonstrated a significant decrease in intracellular ATP content in Pifithrin-α–treated cells (Figure 6K), consistent with diminished mitochondrial function.

Together, these results confirm that p53 is a critical mediator of ISM1-induced cardiomyocyte maturation, and that ISM1 promotes structural and metabolic specialization largely through p53-dependent mechanisms.

## Discussion

In this study, we identify Isthmin-1 (ISM1) as a previously unrecognized regulator of cardiomyocyte maturation. Through cross-species transcriptomic analyses, in vitro functional assays, and in vivo genetic models, we demonstrate that ISM1 enhances sarcomere organization, mitochondrial metabolism, and calcium-handling competence. Mechanistically, ISM1 activates the p53 signaling pathway and directly interacts with p53, enhancing its stability, nuclear translocation, and transcriptional activity to promotes structural and metabolic specialization.

Among the three genes commonly upregulated in both human and mouse datasets—ISM1, ANKRD1, and HBEGF—ISM1 exhibited the highest log₂ fold change and was therefore selected for mechanistic exploration. While ANKRD1 and HBEGF are well-established markers of mechanical stress and hypertrophic signaling^20, 21^, their induction typically reflects secondary adaptive responses to cardiac injury rather than direct regulation of maturation. In contrast, ISM1 expression increased in both adult versus fetal hearts and pathological hypertrophy, suggesting a dual role in promoting maturation and protecting against de-maturation.

As a secreted protein previously linked to angiogenesis and metabolic regulation^22, 23^, ISM1 likely serves as a molecular bridge between extracellular cues and intracellular energy remodeling. In our iPSC-CM system, ISM1 overexpression enhanced structural alignment, oxidative metabolism, and excitation–contraction coupling, while ISM1 knockdown produced the opposite effect. Together, these results establish ISM1 as a maturation factor that facilitates the transition of cardiomyocytes from a glycolytic, proliferative state toward an oxidative, functionally specialized phenotype.

Cardiac hypertrophy and heart failure are characterized by a reversion to fetal gene expression, impaired oxidative metabolism, and disrupted calcium handling^10–13^. Consistent with this paradigm, our findings suggest that ISM1 may suppress fetal gene reactivation and preserve cardiomyocyte maturity under stress, thereby mitigating progression toward heart failure. The precise effects will be further explored in our future studies.

Our RNA-seq analyses revealed that ISM1 overexpression induces global transcriptional reprogramming, including upregulation of catabolic and lipid metabolic pathways and suppression of cell-cycle and proliferative genes—a hallmark of maturation. Among the enriched pathways, p53 signaling was the most prominently activated. p53 is a master regulator of mitochondrial biogenesis, oxidative phosphorylation, and cell-cycle exit, processes crucial for cardiomyocyte maturation^24–26^.

Consistently, ISM1 Co-IP/MS identified p53 itself and 15 proteins known to activate p53 signaling, positioning ISM1 within an upstream regulatory hub of this pathway. Mechanistically, ISM1 directly binds to p53, stabilizing it, delaying its degradation, and promoting its nuclear localization. This stabilization enhances the transcription of key p53 targets such as *CDKN1A, TIGAR*, and *SCO2*, which regulate energy metabolism, redox homeostasis, and mitochondrial respiration. Thus, ISM1 strengthens the p53-driven transcriptional network that underpins cardiomyocyte metabolic and structural maturation.

While this study defines the ISM1–p53 axis as a critical regulator of cardiomyocyte maturation, several questions remain. The upstream signals that govern ISM1 expression under physiological and pathological conditions are unknown. Given ISM1’s secretory nature, it may also mediate intercellular crosstalk between cardiomyocytes, fibroblasts, and endothelial cells, influencing broader remodeling networks. Future work using conditional knockout models and 3D human cardiac organoids will be instrumental in defining ISM1’s spatiotemporal roles in cardiac development, adaptation, and aging.

### Conclusion

In this work, we identify Isthmin-1 (ISM1) as a previously unrecognized and essential regulator of cardiomyocyte maturation. Through integrated cross-species transcriptomic analysis, functional assays in human iPSC-CMs, and mechanistic interrogation of transcriptional networks, we demonstrate that ISM1 promotes structural alignment, mitochondrial oxidative capacity, and calcium-handling specialization—core features of mature cardiomyocytes. Loss of ISM1 leads to global immaturity, highlighting its requirement for achieving a fully differentiated state.

Mechanistically, ISM1 exerts its effects by stabilizing and activating the p53 signaling axis, enhancing p53 nuclear localization and transcription of metabolic and cell-cycle regulatory genes. This ISM1–p53 partnership coordinates mitochondrial biogenesis, oxidative metabolism, and proliferative arrest, thereby driving the maturation program. The identification of p53 as a downstream effector places ISM1 at a central node linking extracellular cues with nuclear transcriptional machinery.

Beyond developmental maturation, our findings suggest that ISM1 may also help preserve cardiomyocyte maturity under pathological stress, given its upregulation during hypertrophy and its ability to suppress fetal gene reactivation in vitro. Whether ISM1 can mitigate de-maturation and protect against heart failure in vivo warrants further investigation.

Altogether, this study establishes ISM1 as a critical upstream regulator of cardiomyocyte maturation and a potential therapeutic target for enhancing iPSC-CM–based cardiac regeneration and combating pathological remodeling.

## Data Availability

The data underlying this article are available in the GEO Database and can be accessed with GSE314323.

## Supplemental Material

Supplemental Methods

Tables S1

Figure S1-S3

